# Regeneration-Associated Cells Improve Recovery from Myocardial Infarction through Enhanced Vasculogenesis, Anti-inflammation, and Cardiomyogenesis

**DOI:** 10.1101/396101

**Authors:** Amankeldi A. Salybekov, Akira T. Kawaguchi, Haruchika Masuda, Kosit Vorateera, Chisa Okada, Takayuki Asahara

**Author notes:** Correspondence to: Takayuki Asahara, MD, Ph.D., Department of Regenerative Medicine Science, Tokai University School of Medicine, 143 Shimokasuya, Isehara, Kanagawa 259-1193, Japan.

## Abstract

**Background:** Considering the impaired function of regenerative cells in patients with comorbidities and associated risk factors, cell therapy to enhance the regenerative microenvironment was designed using regeneration-associated cells (RACs), including endothelial progenitor cells (EPCs) and anti-inflammatory cells.

**Methods:** RACs were prepared by quality and quantity control culture of blood mononuclear cells (QQMNCs). The peripheral blood mononuclear cells (PBMNCs) were isolated from Lewis rats and conditioned for 5 days using a medium containing stem cell factors, thrombopoietin, Flt-3 ligand, vascular endothelial growth factor, and interleukin-6 to generate QQMNCs.

**Results:** *In vitro* EPC colony forming assays demonstrated a 5.3-fold increase in the definitive colony-forming EPCs and vasculogenic EPCs, in comparison to naïve PBMNCs. Flow cytometry analysis revealed that QQMNCs were enriched with RACs, such as EPCs (28.9-fold, P<0.0019) and M2 macrophages (160.3-fold, P<0.0002). Cell transcriptome analysis revealed that angiogenesis (*angpt1, angpt2,* and *vegfb*), stem/progenitor (*c-kit* and *sca-1*) and anti-inflammation related (*arg-1, erg-2, tgfb,* and *foxp3*) genes were highly expressed in QQMNCs. For *in vivo* cell transplantation experiments, 1×10^5^ cells were administered via the tail vein into syngeneic rat models of myocardial infarction (MI). Echocardiographic data showed that QQMNCs-transplanted group (QQ-Tx) preserved cardiac function and fraction shortening (45.5±4.6%) at 28-days post-MI in comparison with PBMNCs-transplanted (PB-Tx) (30.9±6.4%, P<0.0001) and Control (32.2±7.7%, P<0.0008) groups. Histological analysis revealed that QQ-Tx showed enhanced angiogenesis and reduced interstitial left ventricular fibrosis, along with a decrease in neutrophils and an increase in M2 macrophages in the acute phase of MI. Cell tracing studies revealed that intravenously administered QQMNCs preferentially homed to ischemic tissues via blood circulation, while PBMNCs did not. QQ-Tx showed markedly upregulated early cardiac transcriptional cofactors (*Nkx2-5,* 29.8-fold, and *Gata-4*, 5.2- fold) as well as c-kit (4.5-fold) while these markers were downregulated in PB-Tx. In QQ-Tx animals, *de novo* blood vessels formed a “Biological Bypass” as observed macroscopically and microscopically, while PB-Tx and Control-Tx groups never developed epicardial blood vessels but showed severe fibrotic adhesion to the surrounding tissues.

**Conclusion:** QQMNCs, as RACs derived from rat PBMNCs, conferred potent angiogenic and anti-inflammatory properties to the regenerative microenvironment, enhancing myocardiogenesis and functional recovery of rat MI hearts.

## INTRODUCTION

Despite improved pharmacological and surgical interventions, ischemic heart disease (IHD) is the leading ause of premature mortality; since the year 2006, IHD-related mortality has increased by 19% worldwide[1]. Two decades have passed since the discovery of endothelial progenitor cells (EPCs)[2] and several studies have concluded that in addition to cellular replacement of myocardial loss, EPCs of the hematopoietic stem cell (HSC) line secrete paracrine factors which play an essential role in cell to cell communication and the resolution of inflammation and subsequent recovery[3-5]. These paracrine factors can be released from transplanted cells as proteins or extracellular vesicle cargos, along with non-coding single strand miRNAs, a promising therapeutic tool[6].

EPCs are extremely rare in the adult peripheral blood (approximately 0.005 %), and the paucity of these progenitor cells has hampered the collection of adequate cell numbers for stem cell-based therapy[7, 8]. To this end, several granulocyte-colony stimulating factor (G-CSF)-mobilized peripheral blood (PB) CD34+ cell or mononuclear cell (PBMNCs) -based clinical studies have been conducted and modest outcomes obtained[9, 10]. The majority of patients with risk factors, such as smoking[11], aging[12], and hypercholesterinemia[13], and comorbidities, such as arterial hypertension, obesity, and atherosclerosis, present with chronic excessive secretion of inflammatory cytokines, such as IL-6, IL1b, and TNFa, which leads to impairment in the function of regeneration-associated blood cells, including EPCs[14],[15]. In addition, the aforementioned metabolic inflammatory diseases, along with diabetes, are associated with poorer mobilization of EPCs in patients who received G-CSF[16, 17]. They are also involved in cross-talk with bone marrow (BM) or PB-derived MNCs composed of various hematopoietic cell lines used in transplantation after myocardial infarction (MI), increasing the complexity of the disease[18, 19].

Due to these additional complications in the patients with comorbidities or risk factors, the quantity and quality controlled (QQ) culture technique has been proposed to increase regeneration-associated cells (EPCs, and anti-inflammatory macrophages, and T cells) for cardiovascular stem cell therapy[20, 21]. Initially, the QQ-culture method was developed to increase the quality and quantity of vasculogenic EPCs [20]. Under QQ incubation, naïve PB pro-inflammatory (monocytes and macrophages type 1 (M1*ϕ*)) cells convert to anti-inflammatory phenotypes (macrophage type 2 (M2*ϕ*)), and modulate immunity by decreasing cytotoxic T cells and natural killer (NK) cells, consequently increasing the immune-tolerance T-helper subset, regulatory T (Treg) cells, leading to extensive tissue regeneration.

In this present study, we explored whether (1) rat QQMNC therapy promotes angiogenesis, (2) anti-inflammatory effect, and (3) *de novo* cardiomyogenesis induction, and (4) subsequently leads to a reduction in fibrosis and (5) improvement of cardiac function after the onset of MI. The findings of this study would aid the development of QQMNCs as a therapeutic agent for MI and other ischemic diseases.

## MATERIALS AND METHODS

All studies were performed with the approval of the national and institutional ethics committees. The Tokai School of Medicine Animal Care and Use Committee gave local approval for these studies, based on Guide for the Care and Use of Laboratory Animals (National Research Council). A total of 120 rats were used.

### PBMNC isolation and QQ Culture

The PBMNCs were collected after anesthesia from the abdominal aorta using a 10-ml syringe containing heparin (500 IU), and MNCs were isolated by density gradient centrifugation using the Lymphocyte Separation Solution (Histopaque, Nakalai tesque, Kyoto, Japan) as reported previously[21]. QQ culture medium of stem Line II (Sigma Aldrich) contained four rat (rat stem cell factor (SCF), vascular endothelial growth factor (VEGF), thrombopoietin (TPO), and IL-6) and one murine (Flt-3 ligand) recombinant proteins (all obtained from Peprotech). Isolated PBMNCs were cultured for five days at a cell density of 2.0×10^6^ /2 mL per well in QQ culture medium (Stem Line II, Sigma Aldrich) in 6-well Primaria plate (BD Falcon). All the essential materials for QQ culture are given in the Table in **S1 Table**.

### EPC Colony Forming Assay

Freshly isolated PBMNCs and post QQ cultured cells were seeded at a density of 1.5×10^5^ cells per 35-mm Primaria dish (BD Falcon), in semisolid culture medium, as described previously[20]. Eight to ten days after seeding the cells, the number of adherent primitive EPC colony-forming units (pEPC-CFUs) and definitive colony-forming units (dEPC-CFUs) were counted separately using a light microscope (Eclipse TE300; Nikon).

### Flow Cytometry

PBMNCs and QQMNCs were suspended in 2 mmol/L EDTA/0.2% BSA/PBS buffer (FACS buffer) (4×10^5^ cells/200 μL buffer) and 45 μL of the cell suspension was dispensed into four tubes containing a staining antibody (volume and dilutions as per the manufacturer’s protocol) and incubated at 4°C for 30 min. Subsequently, cells were washed twice with 1 mL of 2 mmol/L EDTA/0.2% BSA/PBS buffer. Binding of antibodies to surface markers was carried out in 500 μL fixation buffer (BioLegend#420801) at room temperature for 20 min, following which cells were washed twice with FACS buffer. Then, 500 μL of permeabilization buffer (BioLegend #1214114) was added to the tubes with cells and incubated at room temperature for 15 min, before intracellularly staining cells with an anti-CD68-FITC antibody (BIO-RAD). Finally, cells were washed twice with the FACS buffer and data were acquired on a FACS Verse (BD Biosciences) and analyzed with FlowJo software, version 10.2. All the fluorophore-labeled monoclonal antibodies used are detailed in the Table in **S2 Table**.

### Real-Time Quantitative Polymerase Chain Reaction

The PBMNCs and QQMNCs were collected and stored at −80°C in TRIzol reagent (Invitrogen) for subsequent total RNA isolation. After adequate anesthesia, rats were sacrificed 6 days after onset of MI. Heart tissue was excised and perfused with 25 mL of cold PBS then the infarcted area of the left ventricle (LV) was cut and incubated in RNAlater (Sigma) at 4°C overnight. Immediately after homogenization of heart tissue using 1 mL TRIzol (Invitrogen), genomic DNA was digested by DNase I (Invitrogen) at 37°C for 15 min and the total RNA was purified by phenol extraction and ethanol precipitation. Two micrograms of purified total RNA was reverse transcribed to cDNA using the High Capacity cDNA Reverse Transcription kit (Applied Biosystems). The cDNA mixture was diluted 20 to 160-fold with Milli-Q water (Millipore Corporation, Billerica, MA) and the SYBR green master mix (Applied Biosystems) was added according to the manufacturer’s protocol. The relative mRNA expression was calculated by Livak or ΔΔCt method with normalization to the rat 18S rRNA according to the MIQE guideline[22]. The forward and reverse primer pairs used are listed in the Table in **S3 Table**.

### Rat Myocardial Infarction Induction and Cell Transplantation

Male Lewis rats 6 to 10 weeks of age, weighing 150-250 g (Charles River Laboratories Japan, Inc., Tokyo) were used. For cell tracing experiments, male *Lew-CAG-eGFP* transgenic rats were obtained from The National BioResource Project for the Rat in Japan. To induce MI, animals were anesthetized with 2-4% sevoflurane (Maruishi Pharmaceutical Co., Ltd. Japan), after which animals were orally intubated with a 14G intravenous catheter and respired using rodent ventilator at 10mL/kg, 60 times per min (UGO Basile S.R.I., Italy). After left-sided thoracotomy, the left anterior descending (LAD) artery was ligated with Monoplane 6-0 sutures (Ethicon Inc., USA), as described previously[23]. MI induction was confirmed by observation of blanching and dyskinesia of the anterior wall distal to the suture. The thorax muscles and skin were closed using 4-0 nylon and 3-0 silk, respectively. At day 3 after MI onset, animals underwent cell transplantation with conditioned QQMNCs (QQ-Tx group) or PBMNCs (PB-Tx group), at a dose of 1×10^5^ cells, and the control group animals received RPMI 1640 (GIBCO) medium (Control group) via the tail vein, using a 24G angiocatheter (Terumo).

### Echocardiography

Rat echocardiography (EchoCG) was performed under anesthesia with sevoflurane 2.0%, using an ALOKA ProSound SSD 4000 ultrasound device with a recorder. In this study, all the rats underwent baseline transthoracic echo-Doppler and rats surviving after MI induction were followed up with serial examinations at 1, 2, 3 and 4 weeks post-MI. A two-dimensional short-axis view of the LV was obtained at the level of the papillary muscle as described before[24]. The following formulae were used to calculate ejection fraction (EF) = 100 * (end diastolic volume (EDV) - end systolic volume (ESV)) / (EDV) and fractioning shortening (FS) = 100 * ((LVIDd-LVIDs) /(LVIDd)), accordingly[23, 24]. Cardiac catheterization was performed using a microtip catheter (Miller Instruments Inc., Houston, USA).

### Histological Analysis

Heart tissue washed with heparinized PBS solution (1000 U/500 mL) and was fixed with 4% paraformaldehyde (PFA), and the LV was weighed while filled with PBS as the pure condition, to define LV volume capacity. Then heart tissue was fixed in 4% PFA overnight, embedded in paraffin, and cut into 4-mm-thick sections. The heart tissues of the eGFP positive QQMNC and PBMNC transplanted groups were fixed in 4% PFA and incubated in graded sucrose (5, 10, 15 and 20%, for 2 hours each) and then left overnight in 20% sucrose. Samples were embedded with O.C.T compound (Sakura Tek) and cut into 7 μm-thick sections. Sections were stained with picrosirius red to determine the size of the infarcted tissue and interstitial fibrosis, as described previously.^24^ These data were evaluated from 3-5 high -power fields per section in a blinded manner using Image J (version 1.51, National Institutes of Health). For evaluation of infarcted tissue microvascular density (MVD), sections were blocked in an avidin and streptavidin complex and incubated with Isolectin B4-FITC (1:100) (Invitrogen), aSMA-Cy3 (1:200) (Sigma), and anti-CD31 (1:100) (#ab64543, Abcam). For detection of tissue inflammation, heat-induced epitope retrieval was performed in sodium citrate buffer for 20 minutes at 98°C to expose target proteins. Then, sections were incubated overnight with the antibodies against iNOS (1:100) (#ab15323, Abcam), myeloperoxidase (1:100) (#ab9535, Abcam) and CD206 (#ab64693, Abcam). To determine the proliferation of cardiomyocytes, which is an index of cardiomyogenesis, BrdU (Sigma) powder was dissolved in PBS and filled in a subcutaneous osmotic pump that administered 75 μg/kg BrdU for one week. Spectral analysis was performed with a Carl Zeiss LSM880 Meta confocal microscope to discriminate auto-fluorescence.

### Statistical Analysis

All values are showed as mean ± SEM. Mann-Whitney U and Kruskal Wallis test were used for 2 and 3 non-parametric groups with Dunn’s multiple comparison tests, respectively. For multiple comparisons between groups at different time points, 2-way ANOVA was applied, followed by Tukey’s post hoc test. All statistical analysis was performed using GraphPad Prism 7.1 (GraphPad Prism Software Inc., San Diego, CA, USA). P<0.05 was considered statistically significant.

## RESULTS

### Increased Colony-Forming EPCs in rat QQMNCs

To evaluate the vasculogenic potential of freshly isolated PBMNCs in comparison to cultured QQMNCs, 1.5×10^5^ cells were seeded on to a methylcellulose coated dish, and at day 8 to 10 post-seeding, pEPC-CFUs, and dEPC-CFUs were separately counted **(Fig 1A)**. The total number of EPC-CFU colonies increased in QQMNCs compared to PBMNCs (60±5 vs. 16 ±1.95, *P*<0.0043) **(Fig 1B)**. From the total number of EPC-CFU colonies the proportion of dEPC-CFU and pEPC-CFU statistically significantly higher after QQMNC conditioning (*P*<0.0001 and *P*<0.007), but not in naïve PBMNC **(Fig 1B)**. Furthermore, colonies in QQMNCs were 5.3-fold enriched for dEPC-CFUs in comparison with those of PBMNCs. The differentiation ratios of dEPC-CFUs were 75% and 62.5% for QQMNCs and PBMNCs, respectively **(Fig 1C)**. The dEPC-CFUs were composed of large spindle-shaped cells with strong vessel-like tube formation capability *in vitro* **(Fig 1D),** whereas immature pEPC-CFU were cells unable to form the tube-like structure.

**Fig 1.**
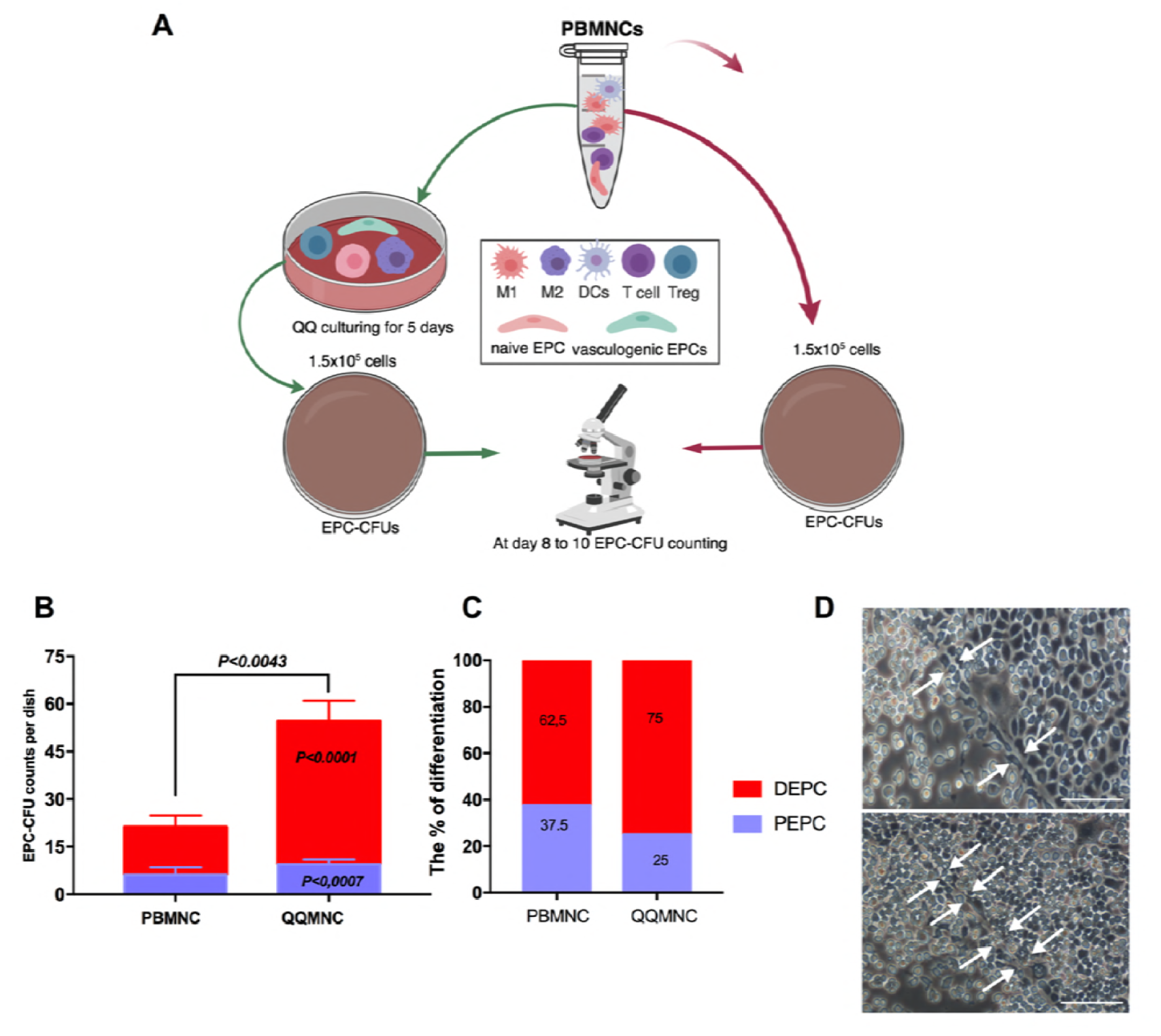
QQ Culture Conditioning Increased Definitive EPC-CFU *in vitro.* **A)** Schematic flowchart of the *in vitro* protocol**. B)** Comparison of EPC-CFU frequency between freshly isolated PBMNCs vs. QQMNCs showed that EPC-CFU counts were higher in QQMNCs **C**) Comparison of the number of EPC colonies (initially seeded cells = 1.5×10^5^ cells/well) with and without QQ culturing, showed that DEPC colony number was increased in QQMNCs **D**) Definitive EPC colonies showed “tube-like” structures (indicated by white arrows) *in vitro*. Statistical significance was determined using Mann-Whitney test. Results are presented as mean ± SEM. Abbreviations: EPC-CFU, endothelial progenitor cells colony-forming units; PBMNCs, peripheral blood mononuclear cells; QQMNCs quality and quantity control culture of mononuclear cells; DEPC, definitive EPC; PEPC, primitive EPC.

Altogether, these data show that QQMNCs increased the number of dEPC-CFUs, which contain further differentiating EPC cells responsible for vasculogenesis.

### Rat QQMNC Expressional Profiles as Regeneration-Associated Cells

After culturing PBMNCs in QQ controlled cocktails, lymphocyte-sized cells (A), monocyte-sized cells, (B) and macrophage-size cells (C) were separately gated from total live cells and compared with freshly isolated PBMNCs using flow cytometry (**S1A Fig).** The pro-inflammatory phenotype, represented by monocyte\macrophages type 1 (M1*Φ*), was estimated by CD68 positivity or CD11b\c double positivity in the total cell population. Freshly isolated PBMNCs were enriched for M1*Φ*, while QQMNCs showed significantly lower M1*Φ* frequency (CD68+: 20±1.85 % vs 5.5±1.3%, CD11b+\c+: 17±2.01% vs 1.25±0.16 %) **(Fig 2A)**. In contrast, anti-inflammatory and regenerative macrophages type 2 (M2*Φ*, CD163^high^CD11b/c^low^), drastically increased in QQMNCs, with approximately 3% in intermediate stages, whereas they were not detected in PBMNCs (13±1.5% vs. 0.06±0.03%; *P*<0.007) **(Fig 2B and S1B Fig)**. This data indicates that QQ conditioning of PBMNCs enriched M2*Φ* through converting the classical M1*Φ* phenotypes toward the alternative activated M2*Φ* phenotype **(Fig 2A-2C and S1B Fig).** Recently, Jablonski et al. defined novel exclusively expressing M1*Φ* (*CD38*) and M2*Φ (erg2* and *arg1)* macrophages markers; we found that the M2*Φ* marker *arg1* was upregulated 323-fold and *erg2* 18-fold in cells post-QQ conditioning, whereas M1*Φ CD38* and *il1b* were significantly downregulated by 5-fold and 2-fold, respectively (**Fig 2F**). The proportion of CD34+ EPCs was significantly increased in cultured cells compared to freshly isolated PBMNCs (28.9-fold; *P*<0.01). RT-qPCR analysis revealed that pro-angiogenic genes, such as *ang1* (430-fold), *ang2* (10-fold), and *vegfb* (323-fold) were strikingly upregulated, except for *pecam 1* and *igf 1* which were downregulated 7- and 2-fold, respectively **(Fig 2C and 2F)**.

**Fig 2.**
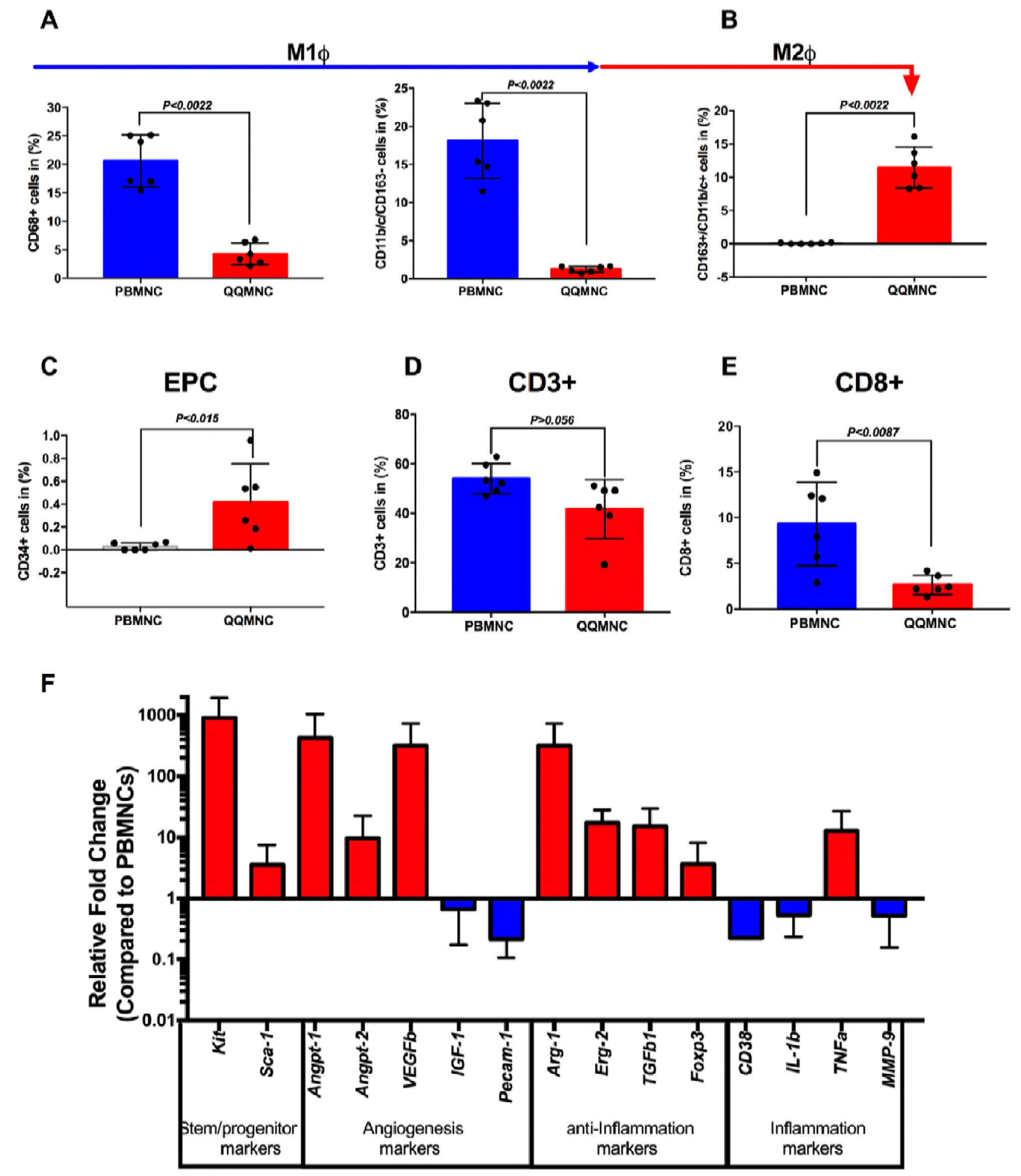
Regeneration-associated Cells Increased post-QQ Culturing. **A)** The flow cytometry analysis revealed that the vast number of PBMNCs were pro-inflammatory monocyte/macrophages (CD68+ and CD11b/c+ /CD163^-^cells). **B)** After QQ culture conditioning, alternatively activated M2*ϕ* percentages were sharply increased. **C**) The proportion of endothelial progenitor cells was higher after QQ culture conditioning. **D)** The number of total T lymphocytes and **E)** cytotoxic T CD8+ lymphocytes, significantly decreased in QQ cultured cells. (**A-E:** n = 6 rats per group) **F**) Relative gene expression profile of QQMNCs compared to PBMNCs (n = 5 rats per group, red bar is upregulated, and blue bar is downregulated, all values are log transformed). Statistical significance was determined using Mann-Whitney test. Results are presented as mean ± SEM.

T-lymphoid cell subsets, particularly cytotoxic T-cells, CD8+ cells, decreased from 10% to 2.3% (*P*<0.0087) of total CD3+ cells **(Fig 2D, 2E and S1C Fig)**. The immunosuppressive T-cell subset regulatory cell marker *foxp3* gene was 4-fold upregulated in QQMNCs compared to PBMNCs **(Fig 2F)**.

Collectively, flow cytometry and RT-qPCR data demonstrated that QQ incubated PBMNCs retained regeneration associated cells, such as anti-inflammatory cells (M2*Φ* cells, *arg1 and erg2* expressing cells), angiogenic cells (CD34+ cells, M2*Φ* cells, *angpt1, angpt2*, and *vegfb* expressing cells), and immunosuppressive T-cells (*Tgfb* and *Foxp3* expressing cells), and reduced cytotoxic T-cells.

### Effect of QQMNCs on LV Performance and Remodeling

The rat body weight (BW) was measured before and 28 days after surgery. At four weeks, BW in the QQ-Tx group increased to 306.4±6 g, in comparison with the PB-Tx (277.6±6 g, P<0.0001), Control (283±8 g, P<0.002), and Sham (282±8 g, P<0.01) counterparts (**Fig 3A**). Serial echocardiography (EchoCG) measurements were performed in QQ-Tx (n=9), PB-Tx (n=11), Control (n=9), and Sham (n=5) groups. Fractional shortening (FS) in the QQ-Tx group increased by day 28 (45.5±4.6%) post-MI, whereas PB-Tx (30.9±6.4%, P<0.0001) and Control (32.2±7.7%, P<0.0008) groups did not recover (**Fig 3B).** EchoCG parameters, including stroke volume and left ventricular systolic dimension (LVSDs) were significantly increased in QQ-Tx (94.4±3.7 μL and 0.35±0.03 cm) while in PB-Tx (63.6±6.07 μL, P<0.004 and 0.49±0.03 cm, P<0.0002) and Control-Tx (71.1±2.0 μL, P<0.03 and 0.46±0.03 cm, P<0.005), LV functional parameters were not enhanced (**Fig 3C and 3D)**. The 2D Doppler mode analysis demonstrates that mild or moderate grade functional insufficiency of the mitral valve was observed in all EchoCG follow-ups in PB-Tx and Control groups while QQ-Tx preserved mitral regurgitation incidence at 4 weeks **(Fig 3E and S1 Video)**. This finding was supported by the Sirius Red staining results showing that in most PB-Tx and Control animals, the LV anterior lateral and posterior lateral papillary muscles were ruptured after MI **(Fig 3F).** Further histological analysis revealed that the infarcted scar area in PB-Tx and Control groups was extended and wall thinning occurred because of excessive inflammation, while these changes were not observed in the QQ-Tx group; therefore, LV remodeling and thickness is promoted by QQMNC transplantation (**Fig 3F, 3H and 3I**). The interstitial collagen fraction was significantly enhanced in the PB-Tx group. In control group, fibrosis was higher, but not statistically significant in comparison with the QQ-Tx group (**Fig 3G and J).**

**Fig 3.**
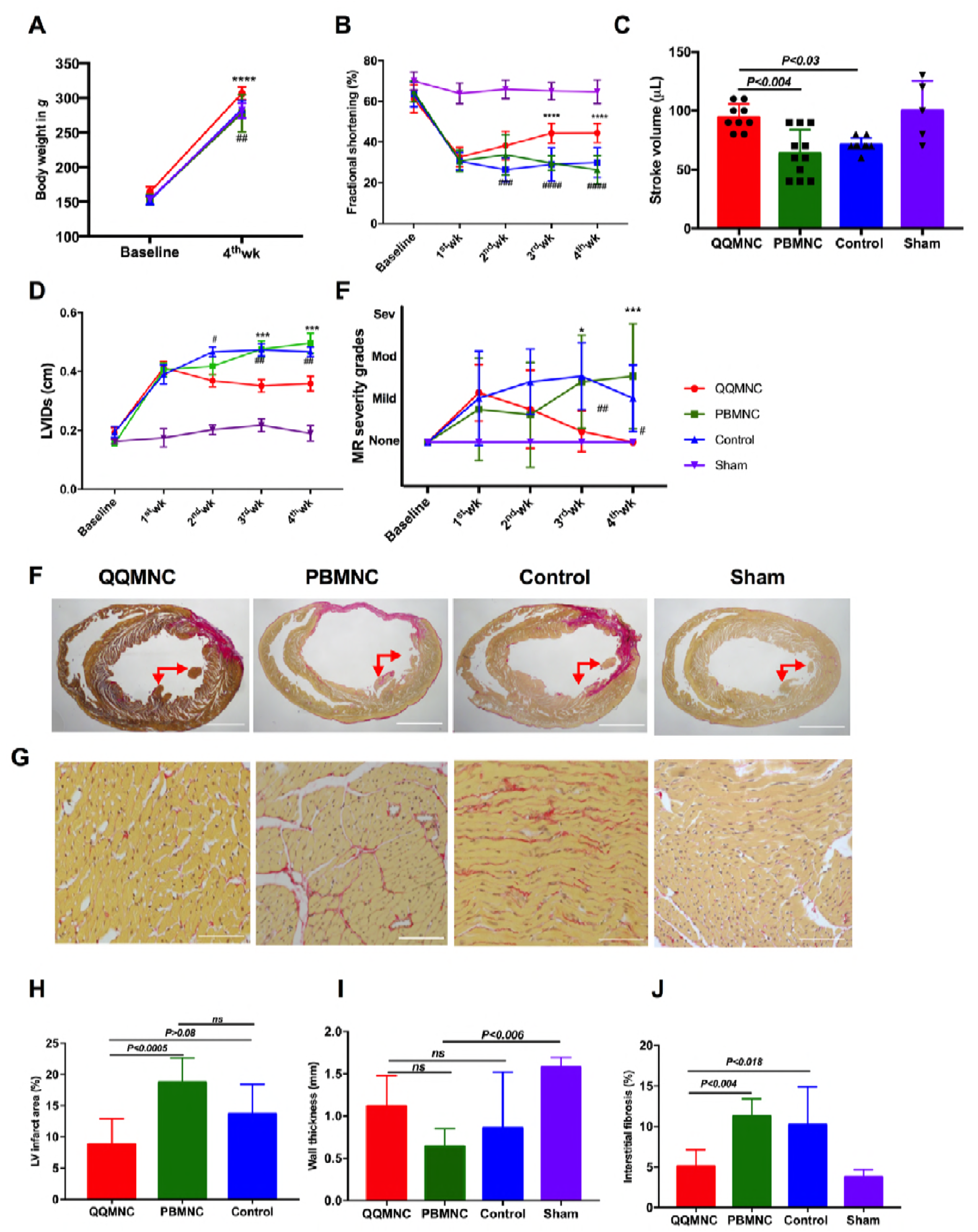
Echocardiographic Parameters Improved after Four Weeks in QQ-Tx Group through Decreasing Fibrosis Composition. **A)** QQ-Tx group animals gained appreciable body weight in 4 weeks. **B)** Fractional shortening was notably increased from the second week after onset of MI in QQ-Tx in comparison with PB-Tx and Control littermates. **C**) Stroke volume was increased and **D)** left ventricular systole dimension was decreased in QQ-Tx group. **E**) Mitral regurgitation events were preserved in QQ-Tx. **F** and **G)** Representative picrosirius red staining of QQ-Tx, PB-Tx, Control, and Sham-operated rat cardiac tissues at day 28 after onset myocardial infarction. **H** and **I**) QQ-Tx group showed preserved left ventricular infarcted area and wall thickness while PB-Tx and Control groups showed extended infarction as well as wall thinning. **J)** QQ-Tx showed reduced interstitial fibrosis. LV-left ventricle. \*\*\*\**P*<0.0001 vs. PBMNC transplanted group; ^#^*P*<0.05; ^###^*P*<0.001; ^*####*^*P*<0.0001 vs Control group. Statistical significance was determined using 2-way ANOVA followed by Tukey’s multiple comparisons test. Results are presented as mean ± SEM.

### QQMNCs Developed Collateral “Biological Bypasses” Into the Infarcted Area

Four weeks later, intraoperationally, we found the “Biological Bypass” phenomena in the QQ-Tx group, where arterial blood was supplied through a bypass distal from LAD ligated areas (**Fig 4A and B)**. This phenomenon was not observed in the PB-Tx and Control groups; rather, they developed a LV aneurysm and severe adhesion to surrounding tissues (thymus and anterior thoracic wall) which complicated the mobilization of the heart (**S2 Fig 2A and 2B**). To determine whether these phenomena are indeed due to vascular bypass, we stained snap-frozen heart tissue slices with Isolectin B4 and αSMA, respectively. The confocal 3-dimension Z-Stack microscopy findings revealed that αSMA stained vessels were greater in number in infarcted tissues, and larger caliber arteria and arterioles supplied the infarcted region, originating from the anterior thoracic wall (probably from aa. intercostales and left internal thoracic aa.) (**Fig 4C and S2 Video**).

**Fig 4.**
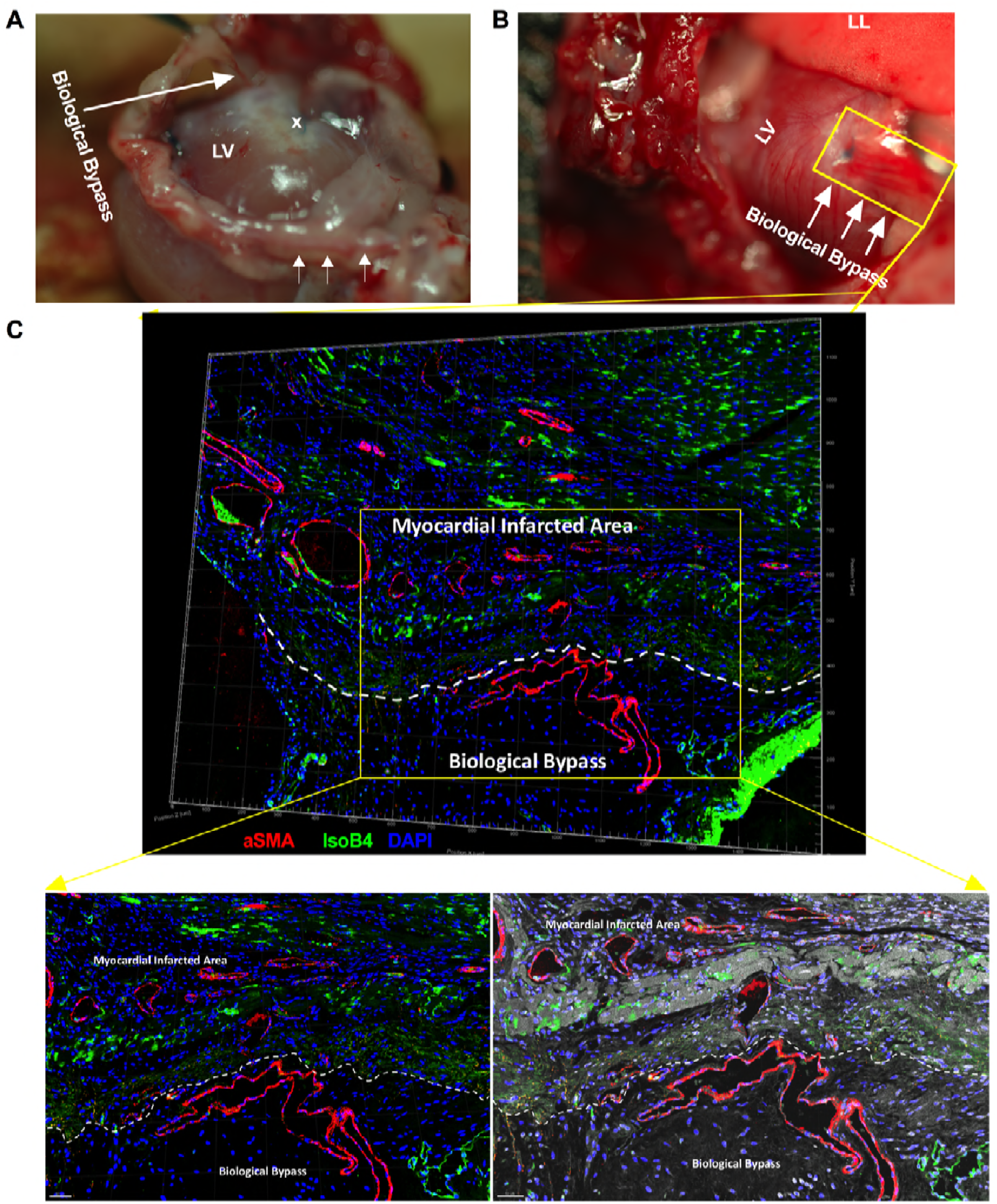
RACs Developed “Biological Bypasses” Into the Infarcted Area. **A)** Intraoperational findings of “Biological Bypass” phenomena in QQ-Tx group are indicated by arrows. The bypass supplied the LV infarcted area. **B)** Several arterioles come from the aa. intercostal distal to the LAD ligation site (indicated by blue monoplane non-absorbable suture) to enhance blood perfusion in myocardial infarcted tissues. Heart tissue was stained (yellow rectangle) with Isolectin B4-FITC to evaluate capillary vessels or with αSMA-Cy3 for pericyte recruited arterioles. **C)** In confocal microscopy, a region of interest (yellow rectangle area) was acquired by tile scan 3-dimension Z-Stack construction, then, all images were fused to obtain a single 3-dimension Z-Stack construction. Myocardial infarcted tissue and biological bypass border area are shown with white dashed line and αSMA stained arterial vessels supplied the myocardial infarcted area. Abbreviations: LV-left ventricle; LL-left lung; x-sutured place. Scale bar: 100 μm (left panel) and 50 μm (right panel).

### Effect of QQMNCs on Angiogenesis in MI

As shown in **Fig 5A**, the microvascular density (MVD) was evaluated by Isolectin B4-FITC staining. MVD was superior in the QQ-Tx group (558±44.4) in comparison with the PB-Tx (260±35.7, P<0.0001) and Control (298±45.5, P<0.0001) groups. To evaluate the functional blood vessel formation^24^, arteriole density was counted per/mm^2^ of the infarcted area and was found to be greater in QQ-Tx than PB-Tx (5.714±2.9, P<0.03) and Control (4.67±2.4, P<0.01) groups **(Fig 5B)**. Confocal microscopy with 3D and Z-stack construction showed that MVD was enhanced in the QQ-Tx compared to PB-Tx and Control groups (**Fig 5C and S3 video**). To investigate whether the transplanted QQMNCs were able to differentiate into endothelial cells forming a structure in the host heart tissue, we used *Lew-CAG-eGFP* transgenic rats. In the immunohistochemistry (IHC) study, transplanted eGFP QQMNCs were seen to express the endothelial marker CD31. Furthermore, increased MVD per/mm^2^ was observed in the host tissue of the QQMNC transplanted group, but not in the PBMNC transplanted group (**Fig 5D)**. The RT-qPCR findings revealed that angiogenesis-related genes such as *angpt1, angpt2,* and *pecam1* were significantly upregulated in the QQ-Tx group (3-fold, 9-fold, and 5-fold, respectively, compared to control), but downregulated in the PB-Tx group (2-fold, 1-fold, and 2-fold, respectively, compared to control) at day 3 post cell transplantation. (**Fig 5E)**.

**Fig 5.**
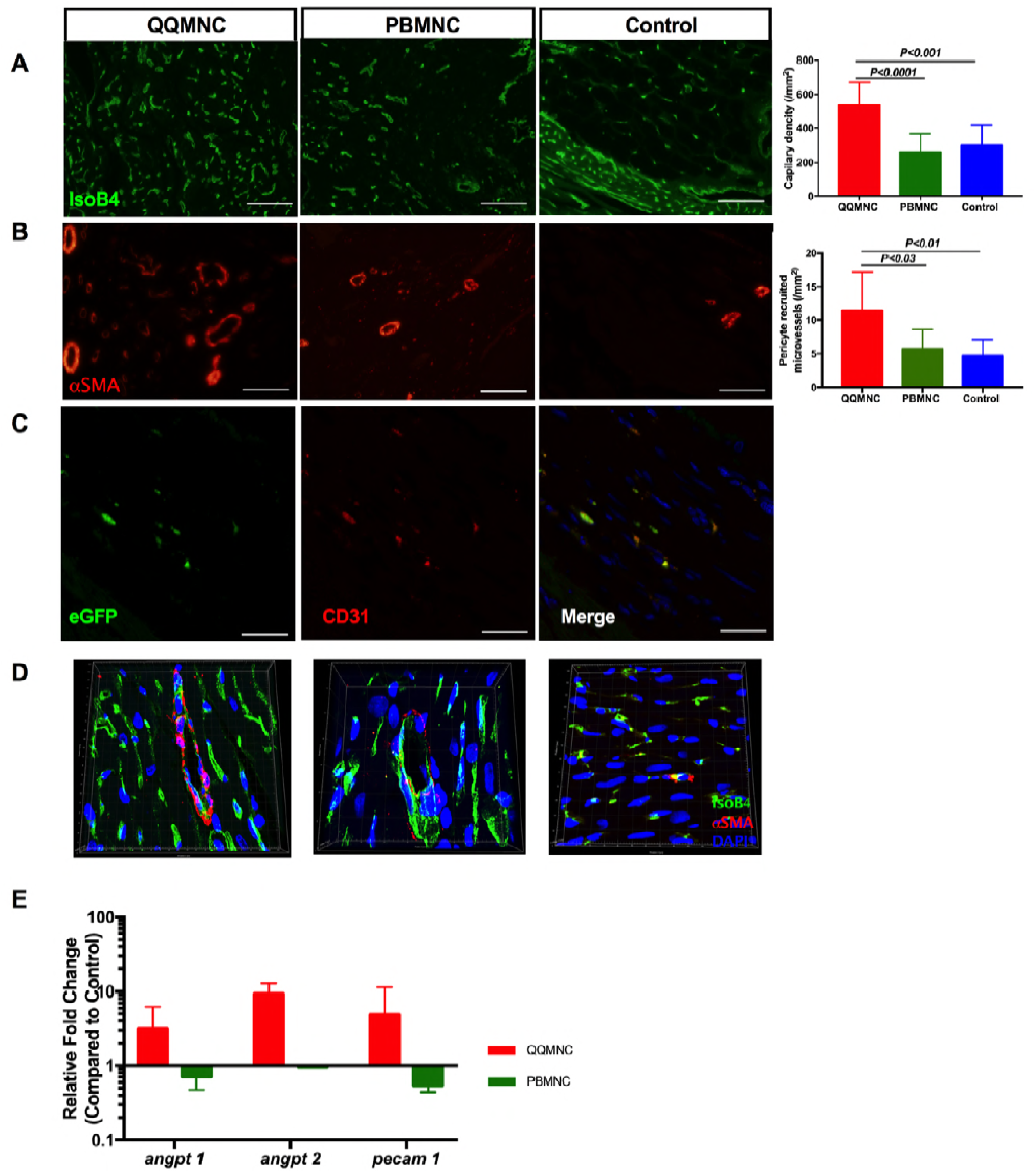
RACs Promoted Angiogenesis and Arteriogenesis in Myocardial Infarcted Area. Six randomly detected fields were evaluated, including border and infarcted areas, with 20x magnification to obtain an average value of vascularization. **A)** QQ-Tx group showed significantly increased functional capillary density and B) αSMA stained arterioles per/mm^2^ whereas PB-Tx and Control groups did not. **C)** QQ conditioned eGFP expressing CD31+ cells extended capillary length with incorporation into host tissues. **D)** Representative Z-stack 3D construction images of Isolectin B4 and αSMA stained infarcted tissues of QQ-Tx, PB-Tx, and Control groups. **E**) At day 6 post-MI, infarcted tissues of QQ-Tx showed elevated angiogenesis-related gene expression (all values are log transformed). Scale bar: 20 μm. Statistical significance was determined using Kruskal-Wallis followed by Dunn’s multiple comparisons test. Results are presented as mean ± SEM.

In summary, QQMNC transplantation promoted vasculogenesis, angiogenesis, and arteriogenesis via accelerating EPC maturation on infarcted tissues.

### Tracing of Transplanted QQMNCs in Infarcted Myocardium and Other Organs

We employed *Lew-CAG-eGFP* transgenic rats to trace transplanted cells. Four weeks after transplantation, animals were sacrificed for immune-histological cell tracing analysis, which revealed that in QQ-Tx animal, eGFP positive cells homed into the infarcted and peri-infarcted areas (**Fig 6A and 6B**), whereas this was not the case in PB-Tx animals (**Fig 6C**). In addition, a significant number of eGFP positive PBMNCs were trapped in parenchymatous organs, such as the spleen and lungs in PB-Tx animals, in comparison with their QQ-Tx littermates (**Fig 6D and 6E, S4 video**).

**Fig 6.**
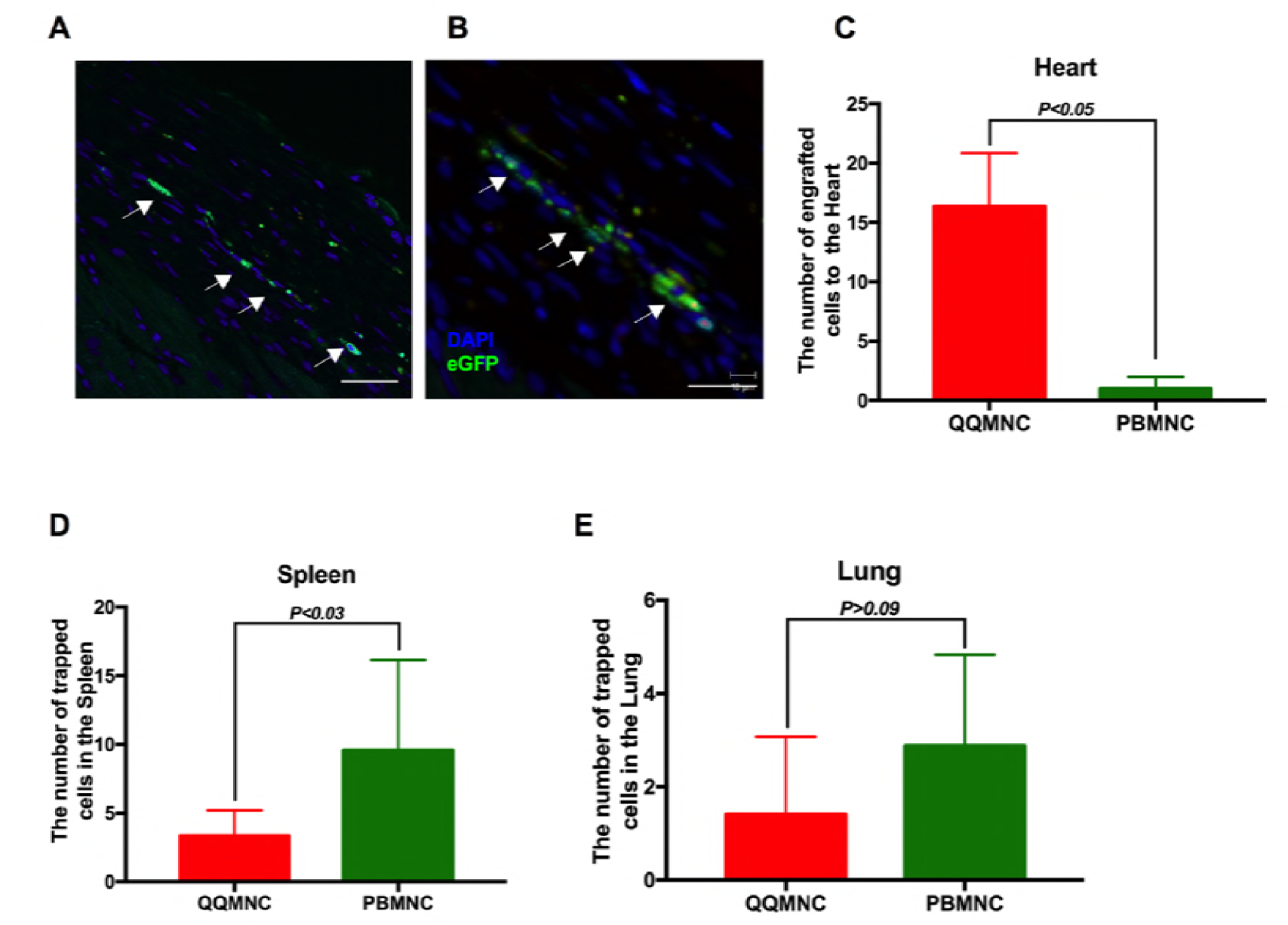
QQMNCs Homed Myocardial Infarcted Tissues. **A)** Representative figures of IHC cell tracking of transplanted eGFP positive QQMNCs recruited into the peri-infarcted and **B)** infarcted area. **C)** A significant number of eGFP positive QQMNCs homed to the infarcted area at 4 weeks in the QQ-Tx but not in the PB-Tx group. **D and E)** Majority of transplanted eGFP PBMNC cells were trapped in the spleen (**D)** and lung tissues (**E)**. Scale bar: 20 μm (A) and 40 μm (B). Statistical significance was determined using Mann-Whitney test. Results are presented as mean ± SEM.

### Effect of Regeneration-Associated Cells on Inflammation

To test whether neutrophil and monocyte/macrophage infiltration is the leading cause of LV remodeling in the acute phase of infarction, we performed IHC staining) at day 6 for inducible nitric oxide synthase (*iNOS*) to evaluate tissue inflammation **(Fig 7A)** and myeloperoxidase (MPO) to detect neutrophils infiltration **(Fig 7B)**. The findings showed that iNOS stained infarcted areas were more widespread in PB-Tx (24.12±1.35%, *P*<0.0007) and Control (27.63±2.63%, *P*<0.0001) groups, than in the QQ-Tx (7.3±0.7%) group, indicating the QQ-Tx animals showed diminished inflammation **(Fig 7A and 7C)**. Accordingly, the MPO expressing neutrophil-rich area was lower in QQ-Tx animals (3.4±1.1%), in comparison with PB-Tx (7.4±0.6%, *P*<0.03) and Control (13.3±3.35%, *P*<0.01) groups **(Fig 7B and 7D)**. The IHC study also demonstrated that infarcted tissue-infiltrated CD206+ anti-inflammatory M2*ϕ* cell number significantly increased in the QQ-Tx group (99 ±11) than in the PB-Tx (34±9, P<0.01) and Control (45±7, P>0.08) groups, respectively **(Fig 7E and 7F)**.

**Fig 7.**
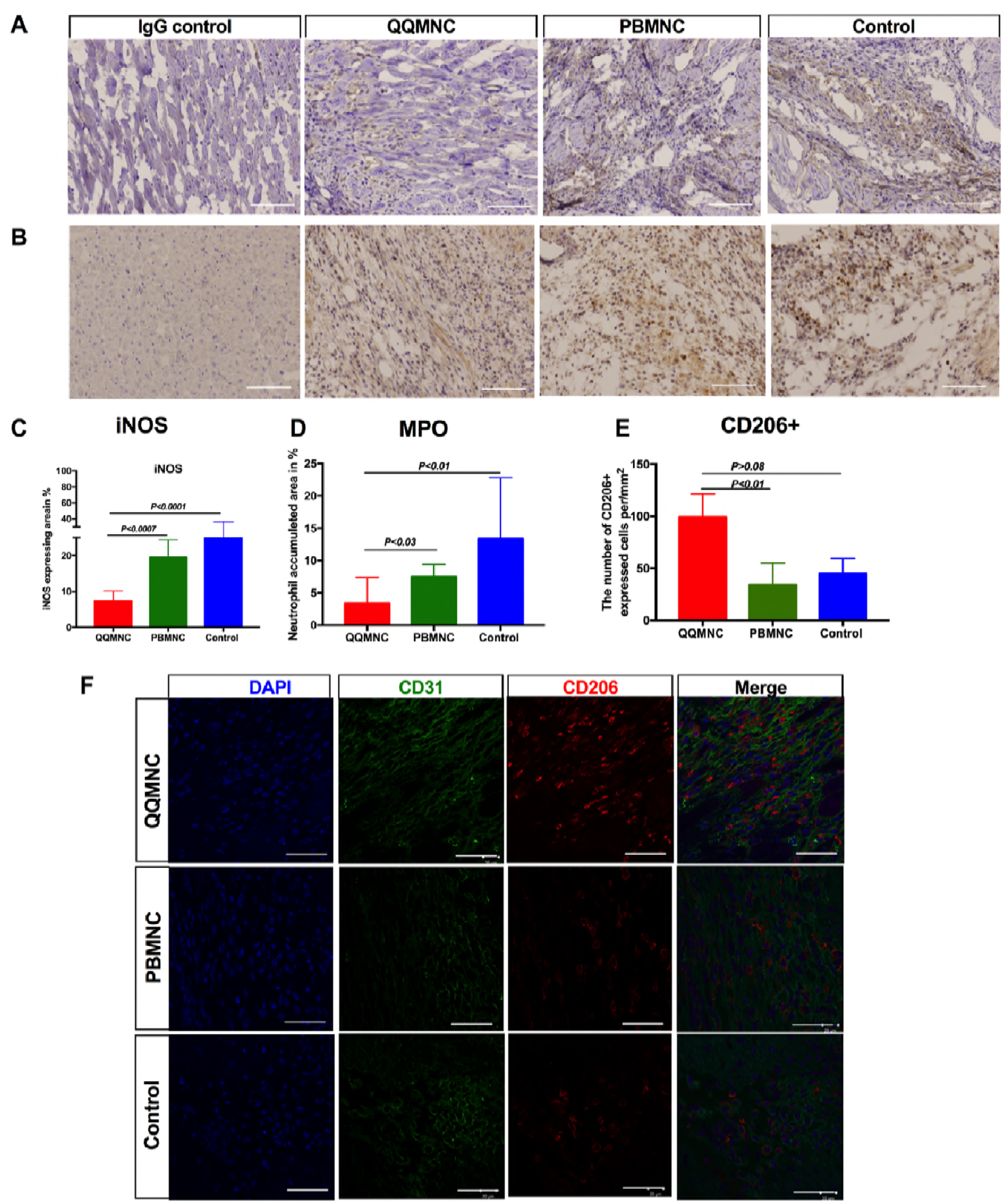
Anti-inflammatory Effect of RACs. **A** and **C**) Immunohistochemical analysis showed that the iNOS expressing infarct area was more extended in PB-Tx and Control groups than in QQ-Tx. **B** and **D**) Neutrophil rich area was significantly lower in QQ-Tx. **E)** The number of infiltrated CD206+ cells after MI were significantly increased in QQ-Tx compared with PB-Tx and Control groups. **F)** Images are showing CD206+ cell recruitment into the infarcted tissue in the acute MI phase. IHC staining was done 6 days after onset of MI. Scale bar: 50 μm for iNOS and MPO, and 20 μm for CD206+. iNOS-inducible nitric oxide synthase; MPO-myeloperoxidase. Statistical significance was determined using Kruskal-Wallis followed by Dunn’s multiple comparisons test. Results are presented as mean ± SEM.

Taken together, the reduced influx of pro-inflammatory cells, neutrophils, and M1*ϕ*, and the shift of phenotypes toward anti-inflammatory M2*ϕ* played a crucial role in the resolution of inflammation and subsequent tissue reparation in QQ-Tx animals.

### Effect of QQMNCs on Cardiomyogenesis

To test whether transplanted cells were able to promote new cardiomyocytes, we performed RT-qPCR analysis to evaluate the relative expression of early cardiomyocyte differentiation cofactors, as reported elsewere[25]. Notably, compared to Control animals, QQ-Tx animals showed marked upregulation of *Nkx2-5* (29.8-fold) and *Gata-4* (5.2-fold), as well as c-kit (4.5-fold), while PB-Tx animals showed downregulated *Gata-4* (1.2-fold) and c-kit (2.5-fold), albeit *Nkx2-5* being 4.7-fold upregulated six days after MI **(Fig 8A)**. Moreover, BrdU incorporated cardiomyocyte nuclei were observed in the QQ-Tx group but not in PB-Tx and Control groups (**Fig 8B**).

**Fig 8.**
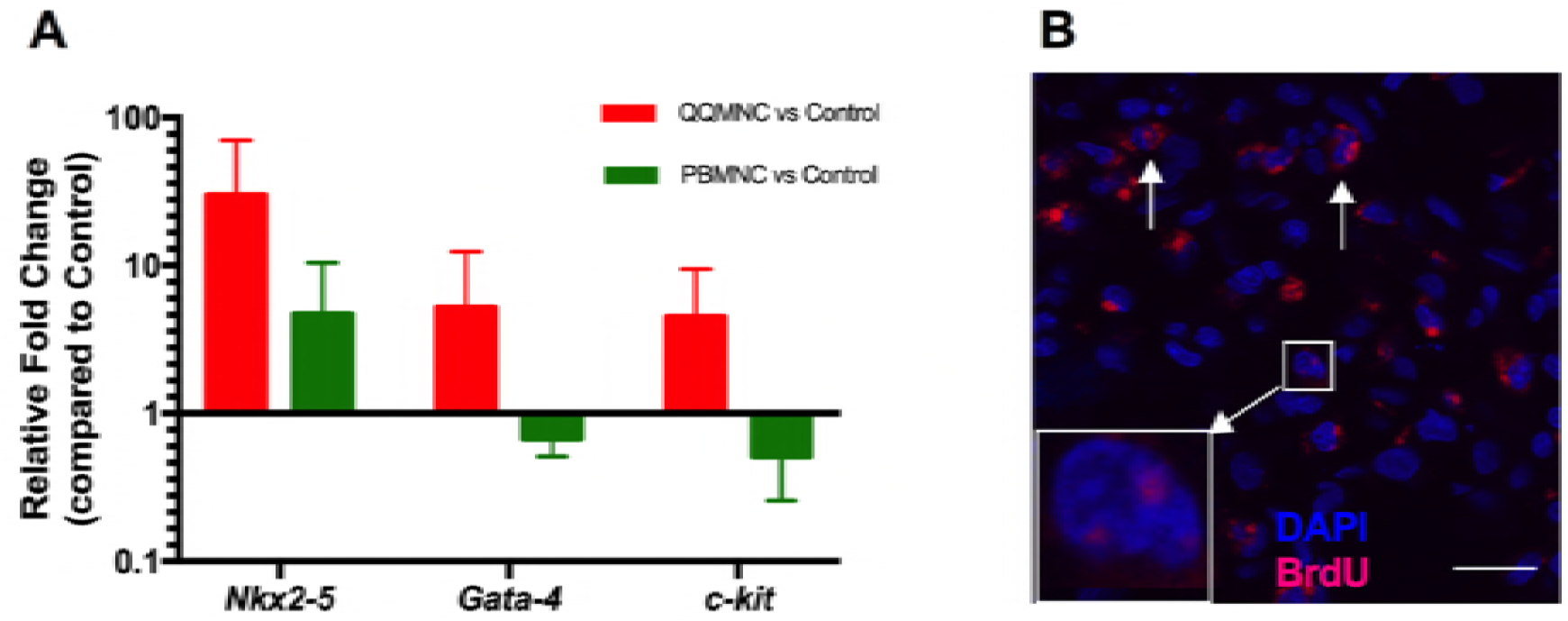
RACs Accelerated Cardiomyogenesis. **A)** Early cardiac differentiation transcription factors were evaluated in infarcted tissues 3 days after cell transplantation (n=5 rats per group, all values are log transformed). **B)** Punctate (square) and diffuse (arrows) localization of bromodeoxyuridine (BrdU) in myocyte nuclei from QQ-Tx group. Scale bar: 20 μm.

## DISCUSSION

In this present study, we have demonstrated that regeneratively conditioned PBMNCs, QQMNCs, were enriched with EPCs and M2 macrophages and increased transplantation efficacy, enabling recovery of ischemic myocardium compared to naïve PBMNCs, by enhancing vasculogenesis, anti-inflammatory properties, and cardiomyogenesis, aiding LV remodeling (**Fig 9**).

**Fig 9.**
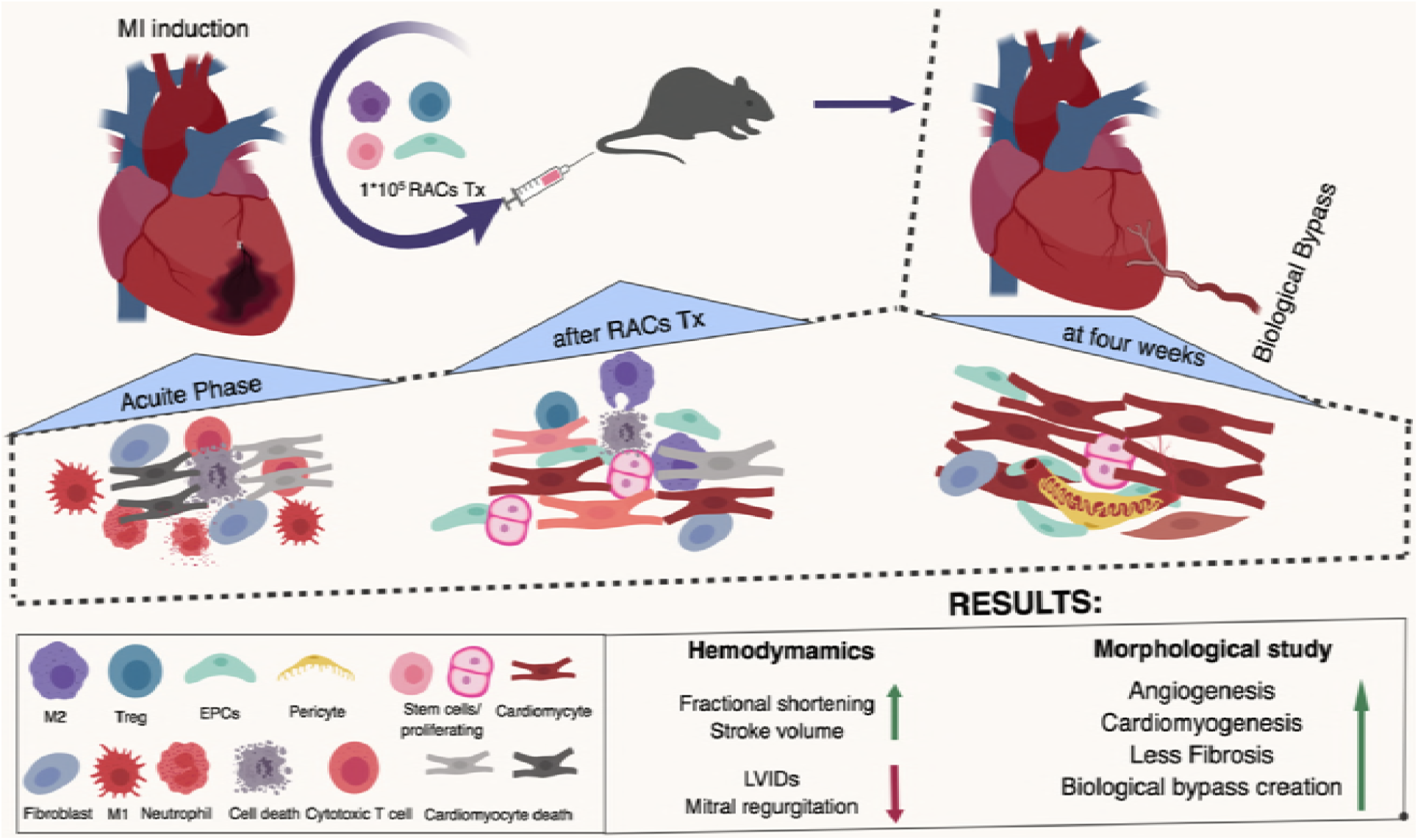
Summary of Our Work.

### QQMNCs as Regeneration-Associated Cells

QQMNCs were developed to enhance the vasculogenic potential of EPCs and facilitate the preparation of monocytes that activate regenerative and anti-inflammatory phenotypes [21]. This method for QQMNC generation is based on an established culture method to increase the quality and quantity of the EPC population, comprising, for example, of CD34+ and CD133+ cells[20]. Interestingly, this vasculogenic signaling condition can educate naïve PBMNCs to not only expand EPCs but also to convert hematopoietic cells into populations responsible for vascular and tissue regeneration^21,29^. With human QQMNCs, definitive colony forming EPCs increased, and macrophages and T lymphocytes became phenotypically polarized into their respective angiogenic, anti-inflammatory, and regenerative subsets. Macrophages transformed from classical M1(CCR2+) cells to alternative M2(CD206+) cells, helper T lymphocytes transformed from Th1 (CD4+/INF-g+/IL4-) to Th2 (CD4+/INF-g-/IL4+) cells, and the angiogenic T cell (CXCR-4+/CD31+/CD3+) and Treg cell (CD4+/CD25+/FoxP3+) populations expanded[21].

In recent years, the interactions between EPCs, monocytes/macrophages, and T lymphocytes have been investigated. EPC angiogenic paracrine factors, such as VEGF, angiopoietins, SEMA3A, SDF1, IL4, etc., are closely related to enhanced anti-inflammatory and immune suppressive phenotypes[26]. IFN-γ produced by Th1 lymphocytes induces monocytes to become classical activated M1 macrophages, while IL-4, IL-13, and IL-10 that are produced by Th2 and Treg lymphocytes induce differentiation of regenerative M2 macrophages. IL-12 and IL-6 produced by M1 macrophages activate Th1 lymphocytes, while IL-10 and TGF-b produced by M2 macrophages induce Th2 and Treg lymphocyte functions[27, 28].

These findings suggest that phenotype changes of naïve PBMNCs in a vasculogenic culture are promoted by crosstalk between EPCs, monocytes, and lymphocytes, and correspond to blood associated cell phenotype transitions in tissue microenvironments in ischemic and regenerative organs, conditioned by angiogenic growth factors, hypoxia, and tissue damage. In this regard, QQMNCs are considered a representative “regeneration-associated cell” (RAC) population for ischemia or regenerative organs. This therapeutic concept was evaluated by several animal experiments, involving hindlimb ischemia, acute ischemic kidney injury, and diabetic wound healing mouse models, which analyzed the potential of QQMNCs to induce recovery organ function[21, 29, 30].

In this study, we first developed a rat QQ culture to produce rat RACs. Rat naïve PBMNCs are composed of 55% lymphocytes (10% cytotoxic T lymphocytes), 20% monocyte/macrophage (approximately 97% is M1*Φ* (CD68+ cells)), and 0.02% CD34+ EPCs. Following QQ conditioning for 5 days, QQMNCs comprise 40% lymphocytes (2.5% cytotoxic T lymphocytes), 15% monocyte/macrophage (75% M2*Φ* (CD163^high^CD11b/c^low^ cells)) and 0.4% CD34+ EPCs. These results strongly suggest that rat QQ culture is optimized to promote RAC development from naïve PBMNCs, in terms of EPC expansion and M2*Φ* conversion, to promote angiogenesis and anti-inflammation

### QQMNC Effect on Vascular Development

Transplanted RACs initiated angiogenesis in host tissues in the same manner as in previously described ischemia disease models[21, 29, 30]. *In vitro,* rat QQMNC EPC-CFUs were increased, mostly with dEPC-CFUs, which are capable of forming “tube-like” structures. Our data are consistent with EPC-CFUs observed in other mouse and human studies[21, 30], which imply that the QQ conditions accelerated the expansion and differentiation of primitive EPC-CFUs (the majority of them in S-phase with high proliferative activity) toward dEPC-CFUs, as reported earlier[8]. Furthermore, pro-angiogenic genes; *angpt-1, angpt-2,* and *vegfb* were markedly upregulated after QQ conditioning. These data indicate the mechanism underlying the enhanced angiogenic quality of QQMNCs.

We observed enhanced new blood vessel formation, in terms of MVD and the number of pericyte recruited arterioles in the infarcted tissues in QQ-Tx rats. eGPF cell tracing experiments disclosed positive RAC-mediated capillary length extension in QQ-Tx host tissues at 4 weeks post-transplantation, but not in PB-Tx tissues.

At day 28, when hearts were exposed during surgery, *de novo* formed, pericardial blood vessels forming a “Biological Bypass” were observed macroscopically in QQ-Tx animals, while PB-Tx and Control-Tx groups developed severe fibrotic adhesion to the surrounding tissues (left lung and thymus). In the rat model of coronary occlusion, c-*kit*-positive cardiac progenitor cells (CPCs) created conductive and intermediate-sized coronary arteries distal to the LAD occlusion and finally the vessels were connected with the primary coronary circulation[31]. Another study demonstrated that when human vascular progenitor cells were transplanted into a dog MI model, large and intermediate caliber newly formed coronary arteries, arterioles, and capillary structures were observed[32]. Ren et al. showed that ERK1/2 and PI3K signaling pathways could stimulate arterial formation and branching in the setting of defective arterial morphogenesis in mice and zebrafish, probably these “Biological Bypasses” could be developed into ischemia tissue by the similar mechanism[33]. The uniqueness of our findings is that the “Biological Bypasses” did not develop from the coronary circulation, but rather arose from either the aa. intercostales or aa. left internal thoracic to enhance infarcted tissue vasculature. The mechanism underlying the development of “Biological Bypasses” needs further elucidation.

### QQMNC Effect on Anti-inflammation

In contrast to naïve PBMNCs, the number of pro-inflammatory cell populations, particularly T cells (cytotoxic CD8+ T cell subset), natural killer cells, and B cells were decreased significantly after QQ conditioning [21],[29, 30].

Data from many previous studies suggest that after ischemic injury to cardiomyocytes, neutrophils first infiltrate the border zone of the at-risk myocardium over 12 h, peaking at day 3, followed immediately by granulocytes, which cause a “cytokine storm.” These early events are followed by pro-inflammatory monocyte/macrophage influx between 5 to 7 days post-ischemic injury, which leads to the spreading of the inflammation process worsening heart function[34, 35]. In our study, the neutrophil-rich area in myocardial infarcted tissue was significantly smaller in QQ-Tx than in PB-Tx and Control groups 3 days after the infarct event. EchoCG and Sirius Red staining revealed that mitral regurgitation incidence was higher in PB-Tx and Control-Tx groups due to papillary muscle rupture and the LV aneurysm. These findings agree with the abundant influx of pro-inflammatory cells, which promote *iNOS* and matrix metalloproteinase activities (*MMP-9*) in the infarcted myocardium, lead to thinning of the LV wall as well as the extension of infarct size[24] by collagen type 1 and 2 depositions, therefore increasing myocardial dysfunction, papillary muscle rupture, and mortality[36, 37].

The PB-Tx animals showed extended LV scarring, interstitial fibrosis, and reduced LV wall thickness in comparison with that observed in a natural course of MI or Control-Tx animals. This can be explained by the fact that 20% of naïve PBMNCs comprise monocytes/ M1*ϕ* cells, increasing the number of pro-inflammatory cells at the site of injury, consequently resulting in the spread of the inflammation process via *CCR2* and *CX3CR1.* On the other hand, in QQ-Tx animals, the infarcted tissues were preserved due to CD206+ cell infiltration [38, 39]. This was supported by RT-qPCR analysis, which demonstrated significant downregulation of distinct M1*ϕ* markers and upregulation of M2ϕ markers in QQMNCs in comparison to in PBMNCs. Pro-regenerative M2*ϕ* secrete several anti-inflammatory cytokines, such as IL-4 and IL-10[40] which activate the T-helper immune-tolerance subset, Treg cells[41, 42]. To confirm the induction of immune-tolerance via Treg function in RAC delivered to tissue, the distinct transcription factor of Treg cells, *foxp3* was upregulated in QQ-cultured MNCs compared to in PBMNCs. The aforementioned immune-tolerance cells play a key role in controlling the inflammation cascade in the acute phase of MI, together with anti-inflammatory M2*ϕ*, to switch on the regenerative phase[43-45]. The *tnfa* was highly upregulated in our study, which may be explained by the initiation of angiogenesis. A previous population-based cancer study also showed that pro-inflammatory cytokines, such as TNFa, IL-6, and VEGF were elevated and promoted the angiogenesis/metastasis process[46].

### QQMNC Effect on Myocardiogenesis

Interestingly, we observed cardiomyogenesis in myocardial ischemic tissue following QQMNC transplantation. Transcriptome analysis of QQMNCs showed high *vegfb* expression. RT-qPCR data of myocardial infarcted tissue also demonstrated that early cardiac differentiation gene expression (*Nkx2-5, Gata-4,* and *c-kit*) was markedly upregulated in QQ-Tx animals on day 6. Emerging evidence suggests that the VEGF-b/SDF-1 axis is important for the mobilization of endogenous CPCs from atrial c-kit+ CPC niches to the infarcted area[47-49]. Perhaps, RACs may promote the recruitment of resident CPCs into the ischemic tissue by regulating the VEGF-b/SDF-1 axis. Another source for CPCs is bone marrow (c-kit+- BMCs). Single-cell-based transcriptional profile analysis of c-kit+-BMCs indicates that genes responsible for engraftment, migration, and differentiation are enriched in cardiomyogenic and vasculogenic c-kit+- BMCs and these cell subtypes not belong to a specific hematopoietic lineage[50]. In a previous clinical study, increased expression of specific cardiac markers (*Nkx2-5, gata-4, and mef2c)* in PBMNCs, and increased number of CD34+/CXCR4+ and CD34+/CD117+ stem cells in PB were observed in ST-elevated MI patients compared to patients with stable angina and healthy Controls[51]. This suggests that QQ-Tx may accelerate endogenous or resident CPC and EPC migration into the ischemic tissue[52], which promotes cardiomyogenesis and angiogenesis.

### QQMNC Recruited into Ischemic Tissue and Effectively Regenerated after MI

One of the exciting findings of this study was the preferential recruitment of intravenously administered QQMNCs into ischemic tissues via blood circulation. The eGFP QQMNC trace experiment revealed that QQMNC retention in organs such as the lung and spleen significantly lower than PBMNC retention, although, physiologically, PBMNCs easily overcome pulmonary passages and enter the systemic circulation. The mechanism of the preferential recruitment of QQMNCs into regenerative tissues is not elucidated in this study, though we can speculate one of the mechanisms for this model. A previous study demonstrated that 7 days of culturing of EPCs increased the expression of CXCR4, the receptor of SDF-1 by approximately 70%, and *in vitro* transmembrane migration assays also revealed the dose-dependent migration of EPCs toward SDF-1[53]. *In vivo*, the SDF-1 expression is modulated by the hypoxia-inducible factor-1 (HIF-1) in endothelial cells which exclusively induces expression of SDF-1 in ischemic tissues in an oxygen tension-dependent manner[54].

In previous clinical trials, a considerable number of autologous PBMNC or bone marrow MNC (BMMNC) (3.0×10^8^ to 5×10^9^) were transplanted into the infarct-related artery through the central lumen of an over-the-wire balloon catheter in MI patients. PBMNC and BMMNC transplantation did not improve regional or global systolic myocardial function during the short-term observation period [55, 56]. Perhaps the implanted PBMNCs or BMMNCs obtained pro-inflammatory phenotypes due to chronic inflammation arising from diseases and risk factors which reduce RACs in quantity and quality. In our study, the small number of QQMNCs administered (1×10^5^ cells, corresponding to ≈ 2.5×10^7^ cells in a human subject with a body weight of 60 kg) proved beneficial for tissue regeneration and functional recovery of the heart from myocardial ischemia.

QQMNCTx significantly increased body weight in comparison with control groups, and even with Sham. The underline mechanism of body weight gain after QQMNCTx needs to further elucidation. The following experiments are necessary to address several key points: (1) finding the optimal dose and (2) timing of transplantation based on the current study, and (3) since QQMNCs demonstrated strong anti-inflammatory and anti-tolerance characteristics, testing in allogeneic experimental transplantation in rat MI models.

## CONCLUSION

In conclusion, the present study demonstrated that *ex-vivo* cultured RACs, which strongly retained several beneficial regenerative characteristics, such as vasculogenesis, immune-suppression, immune-tolerance, and cardiomyogenesis, are a promising therapeutic option for patients with MI and heart failure.

## AUTHOR’S CONTRIBUTIONS

1 - Conception and design, Acquisition of data, Analysis and interpretation of data: S.A.A., A.T.K., T.A., K.V. and C.O., H.M.

2 - Drafting the article, Critical revision of the article: S.A.A., A.T.K., T.A.

3 - Final approval of the version to be published and financial support T.A., A.T.K.

## ACKNOWLEDGMENTS

We thank Ms. Atsuko Sato, Ms. Yoshiko Itoh, Mr. Kazuhiro Yoshida and Mr.Noboru Kawabe for help and assistance with histology and confocal microscopy, Ms. Tomoko Shizuno and Mr. Hiroshi Kamiguchi for help with molecular analysis, Mr. Yusuke Matsumae for help with animal experiments, Dr. Kaori Sekine for help with echocardiography analysis, and Research Support Core Center of Tokai University School of Medicine for outstanding technical support.

